# Spatially resolved tumor-microenvironment architecture defines therapeutic response in unresectable hepatoblastoma

**DOI:** 10.64898/2026.10.10.758234

**Authors:** Zahra Mesrizadeh, Mylarappa Ningappa, Sarangarajan Ranganathan, Kavitha Mukund, Catherine Gestrich, Julia Meade, Judy Hereford Squires, Lori A. Schmitt, George Mazariegos, Marcus Malek, Chethan Ashokkumar, William Andrew MacDonald, Paul Morgan, Shankar Subramaniam, Rakesh Sindhi

## Abstract

**Background:** Hepatoblastoma exhibits variable therapeutic response that is not fully explained by current risk stratification. While prior studies have characterized tumor composition, the role of spatial organization in determining outcome, particularly in unresectable disease requiring transplantation under immunosuppression, remains unclear.

**Methods:** We integrated spatial transcriptomics using the 10x Genomics Visium platform (*n* = 11 tumors) with single-cell spatial molecular imaging using NanoString CosMx (*n* = 3 tumors) on post-chemotherapy explant specimens across clinically defined groups (relapse-prone unresectable, non-relapse unresectable, and resectable hepatoblastoma). Spatial deconvolution, module scoring, spatial co-localization, and cell-cell interaction analyses were used to define region-level transcriptional programs and their underlying cellular organization.

**Results:** We identified three spatial architectures associated with clinical outcome. Relapse-prone tumors exhibited a centralized, TME-dominated organization enriched for progenitor-like, cycling, and stress-responsive tumor cell states co-localized with SPP1+ TAMs and CAFs, with enrichment of hypoxia response, extracellular matrix remodeling, and Midkine (MDK-LRP1) signaling. Non-relapse tumors displayed a distributed architecture with compartmentalized tumor-immune interactions, increased immune engagement, and preserved epithelial differentiation, including inflammatory hepatocyte-like states. Resection samples showed reduced tumor cell state heterogeneity and more uniform TME-immune organization.

**Conclusions**

Integration of regional and single-cell spatial transcriptomics across two platforms reveals that the chemo-remodeled tumor architecture of hepatoblastoma encodes spatially organized tumor cell-TME interactions as central determinants of therapeutic response.

**Translational Relevance:** Current risk stratification in hepatoblastoma — principally based on staging and clinical features — does not reliably predict therapeutic response, particularly in unresectable disease. Our findings identify that spatial tumor-microenvironment (TME) organization is a key determinant of patient outcome. Specifically, our analysis reveals that disease relapse is driven by specialized macrophage-fibroblast signaling networks that promote an immune-cold, therapy-resistant niche. We define a spatially derived gene signature capturing this high-risk TME, which demonstrates strong prognostic value across independent cohorts, highlighting its potential utility as a biomarker for refined patient stratification. These results strongly suggest that incorporating spatial biomarkers into clinical assessment can improve the choice of therapy. Furthermore, the identification of Midkine (MDK) signaling and macrophage-stromal crosstalk as key components of relapse-associated architecture provides a direct rationale for targeting TME communication to combat therapeutic resistance. Together, this work supports a crucial shift from composition-based to spatially informed approaches for risk assessment and therapeutic targeting in hepatoblastoma.

## Introduction

Hepatoblastoma is the most common primary liver malignancy in children, typically arising before the age of four and frequently associated with developmental anomalies (1–3). Although multimodal treatment with chemotherapy and surgical resection have improved survival, patients with unresectable tumors who require liver transplantation remain at risk of rapid and often fatal relapse, as post-transplant immunosuppression compromises tumor immune surveillance (4,5). Current risk stratification frameworks, including the PRETEXT staging system, histological classification, and serum alpha-fetoprotein (AFP) levels, provide important clinical guidance but remain insufficient to fully explain variability in therapeutic response.

Unlike many adult cancers, hepatoblastoma exhibits a relatively low mutational burden (6,7), suggesting that tumor heterogeneity is driven less by genetic alterations and more by dysregulated developmental programs. Indeed, HB tumors recapitulate features of fetal liver development, with distinct epithelial states ranging from differentiated hepatocyte-like populations to progenitor and mesenchymal-like programs. These transcriptional states are governed by key developmental pathways, including Wnt/β-catenin, Notch, and Hedgehog signaling, and are associated with differences in proliferation, differentiation, and clinical outcome (8–12). In parallel, the tumor microenvironment (TME) of hepatoblastoma is increasingly recognized as a critical regulator of tumor progression and therapeutic response (13). Despite the success of immune checkpoint blockade (ICB) therapies in adult cancers, their efficacy in pediatric tumors, including hepatoblastoma, has been limited, potentially due to low tumor mutational burden and reduced T-cell infiltration characteristic of pediatric malignancies (14). These features, combined with the embryonic origin of hepatoblastoma, suggest a fundamentally distinct and potentially immune-restricted tumor microenvironment. While prior studies have begun to characterize hepatoblastoma immune composition using bulk and single-cell approaches (13,15,16), a comprehensive understanding of its spatially organized tumor microenvironment (TME), particularly following chemotherapy, remains lacking.

Here, we integrate region-level spatial transcriptomics with single-cell spatial molecular imaging across clinically defined groups (unresectable (relapse-prone and non-relapse), and resectable hepatoblastoma samples) to define the spatial organization of hepatoblastoma following chemotherapy. By combining Visium-based deconvolution with CosMx single-cell spatial profiling, we characterize how tumor cell states, stromal populations, and immune cells are organized into distinct spatial architectures across clinical outcomes. We show that in unresectable group, relapse-prone tumors are characterized by a macrophage-fibroblast-centered regulatory structure associated with hypoxia-adaptive tumor states and Midkine signaling, whereas non-relapse tumors exhibit distributed tumor-immune interactions and preserved differentiation hierarchies. Together, these findings demonstrate that therapeutic response in unresectable hepatoblastoma is encoded in spatial organization of cell-type and-state composition.

## Methods

### Human Subjects and Clinical Outcomes

All studies were conducted with informed consent and approval from the University of Pittsburgh Institutional Review Board (IRB #0405628). Twelve children with hepatoblastoma (HB) were included in this study, with a median age at liver transplantation (LT) or primary tumor resection of 3.1 years (range 0.99–6.45 years); ten patients were male (Supplementary Table S1). Ten patients underwent liver transplantation for unresectable disease, and two underwent primary tumor resection. all patients received neoadjuvant chemotherapy prior to explanation, and tumor specimens were obtained at the time of transplantation or resection. Primary liver tumor specimens were obtained from patients treated between 2007 and 2019. Among the HB tumors analyzed, two developed post-transplant recurrence (relapse) and died from progressive disease, whereas ten remained non-relapsed at last clinical follow-up (median 121 months) (Supplementary Tables S1 and S2). In the two relapsed patients, relapse occurred at 57 and 61 days after LT, and death following relapse occurred at 15 and 535 days, respectively. PRETEXT stage at diagnosis was available for eleven tumors and ranged from stages I to IV, with one tumor lacking diagnostic imaging at presentation (Supplementary Tables S1 and S3). Relapse status was determined by longitudinal radiologic follow-up and post-treatment histopathologic evaluation. Post-treatment pathologic assessment showed variable responses to chemotherapy, ranging from extensive tumor necrosis to residual viable disease, with diverse histologic and metastatic features observed (Supplementary Tables S2 and S3).

### Spatial Transcriptomics of Hepatoblastoma Tumors Using the Visium Platform

Spatial transcriptomic profiling (ST) was performed on formalin-fixed, paraffin-embedded (FFPE) liver tumor tissue sections from twelve children with hepatoblastoma (Supplementary Table S1) using the Visium Spatial Gene Expression platform and the FFPE Gene Expression Starter Kit for Human transcriptome (Product code#1000334,10x Genomics, Pleasanton, CA). FFPE tissue sections were placed onto the 6.5 mm × 6.5 mm capture area of Visium slides containing approximately 5,000 spatially barcoded spots. Each spot comprised oligonucleotides including a 16-nucleotide spatial barcode for positional identification, a 30-nucleotide poly(dT) sequence for mRNA capture, a 12-nucleotide unique molecular identifier (UMI), and an Illumina Read 1 sequencing adapter. Tissue sections were deparaffinized and stained with hematoxylin and eosin, followed by high-resolution imaging to enable spatial alignment of transcriptomic data. Crosslinks were reversed to release RNA, and transcriptome-wide probe sets were hybridized to target mRNA molecules. Probe pairs bound to RNA were ligated, after which tissues were treated with RNase and permeabilized to release ligated products for capture on the Visium slide. Captured products were extended via a polymerase reaction to incorporate complementary sequences of the spatial barcode and UMI. Spatially barcoded probe products were released and PCR-amplified using sample-specific indexing primers. Final libraries were sequenced to a depth of approximately 25,000 raw reads per capture spot containing tissue using a NovaSeq 6000 system at the UPMC Genome Center. Sequencing data were aligned to histologic images using Space Ranger software (10x Genomics, Pleasanton, CA) and visualized with Loupe Browser. Each capture spot measured 55 μm in diameter with 100 μm center-to-center spacing, corresponding to an estimated average of 5–10 cells per spot, depending on tissue cellularity. Of the twelve tumors processed for Visium analysis, one failed quality-control criteria and was excluded from downstream spatial transcriptomic analyses.

### Single Cell Spatial Molecular Imaging of Hepatoblastoma Tumors Using CosMx SMI

Spatial molecular imaging (SMI) was performed on FFPE liver tumor tissue sections from five children with hepatoblastoma (Supplementary Table S1) using the CosMx™ Spatial Molecular Imager (NanoString Technologies, Bruker; Seattle, WA) and the Human Universal Cell Characterization of 6000 genes panel (part no. 121500041), following the manufacturer’s protocol (MAN-10184-02). Tissue sections were cut at 5 μm thickness, mounted on Superfrost Plus slides, and baked overnight at 60 °C. Sections were deparaffinized, subjected to antigen retrieval at 95–100 °C for 15 minutes, and permeabilized with Proteinase K (3 μg/mL) at 40 °C for 30 minutes. Fiducial markers were applied to facilitate image registration, followed by post-fixation in neutral buffered formalin and blocking with N-hydroxysuccinimide (NHS) acetate. In situ hybridization was performed by overnight incubation at 37 °C with RNA-specific probe sets targeting approximately 6,000 transcripts. After stringent washes, nuclei were stained with DAPI, and cell-segmentation markers including CD298/β2-microglobulin (B2M), pan-cytokeratin (PanCK), and CD45 were applied. Prepared slides were imaged, and iterative cycles of branched fluorescent probe hybridization enabled highly multiplexed RNA detection at single-cell resolution. Images were processed and decoded using the AtoMx™ Spatial Informatics Platform, and the resulting spatial expression data were exported for downstream analysis using open-source bioinformatics tools.

### Visium data preprocessing and quality control

Twelve tissue sections were processed using the Visium spatial transcriptomics platform 10x Genomics, Pleasanton, CA). Downstream analysis of Visium data was performed using the Seurat package (v4.3.0) (17). One sample was excluded due to poor quality, resulting in a final dataset of 11 samples. Each sample was independently normalized using Seurat’s SCTransform (SCT) method (18) and subsequently merged into a combined dataset comprising 16,481 features across 51,016 spots. Quality control filtering was then applied to the merged dataset, retaining spots with a total Unique Molecular Identifier (UMI) count between 500 and 50,000, more than 500 detected genes, and less than 10% hemoglobin gene content. This filtering yielded a final dataset of 16,481 features and 49,505 spatial spots. To identify genes with the highest expression across the spatial transcriptomics dataset, we extracted the raw count matrix from the Spatial assay within the merged Seurat object. Counts were normalized by dividing each gene count by the total number of counts per spot to account for sequencing depth. The top 20 most highly expressed genes were ranked by total normalized expression. The highly abundant liver-specific transcript ALB (albumin) was excluded from downstream analysis to reduce tissue-specific expression bias. After this exclusion, the dataset consisted of 16,480 features and 49,505 spots. Dimensionality reduction and clustering were performed on the SCT-normalized data. Principal component analysis (PCA) was conducted, followed by construction of a shared nearest neighbor (SNN) graph using the top principal components. Louvain-based clustering was applied to identify transcriptionally distinct spatial domains, and Uniform Manifold Approximation and Projection (UMAP) was used for two-dimensional visualization of the clustering results.

### Visium data feature Selection and deconvolution

For deconvolution, STdeconvolve (19) was applied to each of the 11 samples individually. STdeconvolve uses Latent Dirichlet Allocation (LDA) framework to model each spot as a mixture of latent transcriptional programs (“topics”), where each topic represents a gene expression distribution that may correspond to a cell type or cell state. Feature selection was performed by removing genes expressed in fewer than 5% or more than 95% of pixels. Overdispersed genes-defined as genes exhibiting higher variability than expected given their mean expression-were identified using a generalized additive model with five basis functions and an adjusted p-value threshold of 0.05. The 1,000 most significantly overdispersed genes were retained for model fitting. LDA models were trained on each dataset using a range of topic numbers (K) from 2 to 10. For each sample, the model with K = 5 was selected as it achieved the lowest perplexity (a measure of model fit reflecting how well the model predicts unseen data) while avoiding overfitting to rare cell types, defined as deconvolved cell types with mean spot proportions less than 0.05. Cell-type-specific marker genes were identified from the LDA output by calculating log2 fold changes in expression between each deconvolved cell type and all others. Genes with log2 fold change greater than 1 and high expression levels were retained as top markers (Supplementary Table S4). Each deconvolved transcriptional profile, along with its inferred spot-level inferred topic proportions, was mapped back to the merged Seurat object for downstream analysis. Topic assignment was performed using the STdeconvolve method. t-SNE was used to visualize the structure of the deconvolved transcriptional profiles, where visually defined groupings were used to guide topic assignment based on the highest correlation with their transcriptional profiles. These topic assignments were then mapped onto Seurat-defined clusters. Using marker gene profiles from both STdeconvolve and differentially expressed genes (DEGs), each deconvolved cell type was annotated into major biological regions: tumor, stroma, fibrosis, immune mixtures, and hepatocyte. These regional annotations were further validated by a pathologist based on histological inspection of H&E images.

### Visium data enrichment and visualization

Region-specific differential expression gene (DEG) analysis was then performed by comparing each annotated region to all other regions within the same sample. Differentially expressed genes were identified using Seurat’s FindAllMarkers function with the Wilcoxon rank-sum test (parameters: logFC > 0.25, min.pct > 0.25, adjusted p-value < 0.05). Multiple testing correction was performed using the Benjamini–Hochberg false discovery rate (FDR) method. Functional enrichment analysis of DEGs was conducted using the clusterProfiler (20) and Enrichr (21) packages in R. Gene Ontology (GO) analyses for biological processes and molecular functions were performed using the enrichGO function. To explore molecular interactions, a high-confidence human protein-protein interaction (PPI) network (confidence score > 0.85) was obtained from the STRING database v11.5 (22), containing 12,402 proteins and 139,384 interactions. Transcription factor (TF)-target gene relationships from the TRRUST database (23) were integrated with the PPI network to generate a custom TF-PPI network for visualizing regulatory programs associated with region-specific DEGs. All networks were constructed, annotated, and analyzed in Cytoscape (24).

### CosMx data preprocessing and quality control

Single-cell spatial transcriptomic data were generated using CosMx™ Spatial Molecular Imager (NanoString Technologies, Bruker; Seattle, WA) platform. Raw decoded data, including per-cell gene expression matrices, spatial coordinates, and cell-level metadata, were obtained from the AtoMx analysis pipeline (NanoString Technologies, Bruker; Seattle, WA), which performs automated cell segmentation and transcript assignment based on imaging-derived morphology and transcript localization. Downstream analysis of CosMx data was performed using the Seurat package (v5.4) (25). Decoded data were imported into R using NanoStringNCTools package and converted into Seurat objects. Three CosMx datasets were processed and merged for initial quality control and metadata harmonization. Cell-level annotations, including field-of-view (FOV) identifiers and patient labels, were assigned based on NanoString metadata and external sample manifests. Cells lacking patient annotation were excluded before downstream analysis. Quality control was performed using cell-level metrics, including nCount_RNA, nFeature_RNA, and percent.NegPrb. Cells were retained if they satisfied the following criteria: nCount between 500-10,000, nFeature_RNA between 400-5,000, and percent.NegPrb ≤ 10%. Negative control probes and system control probes were separated into independent assays and excluded from downstream expression analysis. This filtering yielded a final dataset of 6,175 features and 348,350 single-cells. To preserve biologically meaningful spatial heterogeneity and avoid cross-sample integration artifacts, all downstream analyses were performed independently for each patient. Cells were analyzed separately for each patient and normalized using SCTransform with 3,000 variable features per sample. For each patient, PCA was performed, and the top 20 principal components were used to construct a shared nearest neighbor graph. Clustering was performed using the Louvain algorithm across multiple resolutions (0.4-0.6), with resolution 0.5 selected based on cluster stability. UMAP was used to visualize patient-specific cellular structure.

### CosMx data cell type annotation

Cell clusters were annotated independently for each patient using a combination of 1) Canonical marker gene expression, 2) differential expression analysis (FindAllMarkers), and 3) spatial localization within tissue. Marker panels included epithelial (e.g., EPCAM, AFP), immune (e.g., PTPRC, CD3D, CD68), stromal (e.g., COL1A1, DCN), and endothelial (e.g., VWF, CDH5). Annotations were refined iteratively to ensure consistency across transcriptional and spatial features.

### CosMx spatial reconstruction

Cell centroid coordinates provided by the CosMx platform were used to reconstruct spatial organization. Coordinates were incorporated into Seurat as field-of-view (FOV) objects, enabling visualization of cell-type distributions within tissue architecture.

### CosMx data differential expression and pathway analysis

Differential gene expression analysis was performed independently for each patient and annotated cell population using Seurat’s FindAllMarkers function. Functional enrichment analysis was conducted using MSigDB (26) gene sets (C5 category) and fGSEA (27), with ranked gene lists defined by signed-log10 adjusted p-values.

### Cell-cell interaction and neighborhood analysis

Intercellular signaling was inferred using the CellChat (28) framework, which models communication probability based on the expression of curated ligand-receptor pairs. For each patient-specific dataset, normalized expression data and annotated cell identities were used as input. Communication probabilities were computed using the default CellChat probabilistic model, which accounts for the average expression of ligands in sender cell populations and receptors in receiver populations. Statistical significance of inferred interactions was assessed through permutation testing, with multiple testing correction performed using Benjamini-Hochberg false discovery rate (FDR) method. High-confidence interactions (adjusted p < 0.01, top 10% interaction strength) were retained for downstream analysis.

To incorporate spatial context, cell-cell proximity relationships were modeled using Giotto (v1.0.4) (29). For each patient, spatial coordinates derived from CosMx cell centroids were used to construct spatial networks. A Delaunay triangulation-based graph was generated to define adjacency relationships between cells, enabling unbiased identification of neighboring cells independent of predefined distance thresholds. This approach captures local tissue architecture while preserving connectivity across irregular spatial distributions. From the Delaunay network, spatial neighborhoods were defined as first-order adjacent cells (direct graph neighbors). Cell-type-specific neighborhood enrichment was quantified using Giotto’s cellProximityEnrichment function, which compares observed cell-cell adjacency frequencies against randomized spatial permutations to identify statistically enriched or depleted interactions.

### Integration of Visium and CosMx data

To relate region-level transcriptional programs identified in Visium data to single-cell spatial organization, we employed a marker-guided annotation strategy. Region-specific gene signatures derived from STdeconvolve analysis of Visium data described above were used to define transcriptional programs associated with tumor, TME, and hepatocyte-enriched regions. These marker sets were then evaluated within the CosMx single-cell dataset to identify spatial regions exhibiting concordant transcriptional profiles. Cells in the CosMx dataset were scored using these Visium-derived marker gene set via AddModuleScore function, enabling assignment of region-associated transcriptional states. Following region annotation, the cellular composition and heterogeneity within each region were characterized using CosMx-defined cell types and spatial organization. This approach enabled resolution of the cell populations and interaction networks underlying Visium-defined spatial domains at single-cell resolution.

## Results

### Integration of Visium sequencing and spatial molecular imaging analysis

To characterize spatial organization of post-chemotherapy explant specimens, we performed spatial transcriptomics (ST) analysis using the Visium platform (10x Genomics, Pleasanton, CA) on eleven tumor sections representing distinct clinical groups unresectable (relapse-prone and non-relapse), and resectable hepatoblastoma samples **(Supplementary Table S1)**. Histological annotation of H&E-stained slides delineated distinct tissue regions across section **(Fig. 1A-B** and **Supplementary Table S2)**. Three representative samples, one per clinical group, were selected for CosMx spatial molecular imaging (SMI; NanoString Technologies, Bruker; Seattle, WA) profiling with multiplex panel targeting ∼6000 genes to capture canonical spatial architectures and to resolve the cellular composition underlying Visium-defined regions at single-cell resolution **(Supplementary Table S1)**. Here, cell types refer to canonical lineage-defined populations, whereas cell states denote context-dependent transcriptional programs within these populations, such as activation, stress, or differentiation states.

**Fig. 1.**
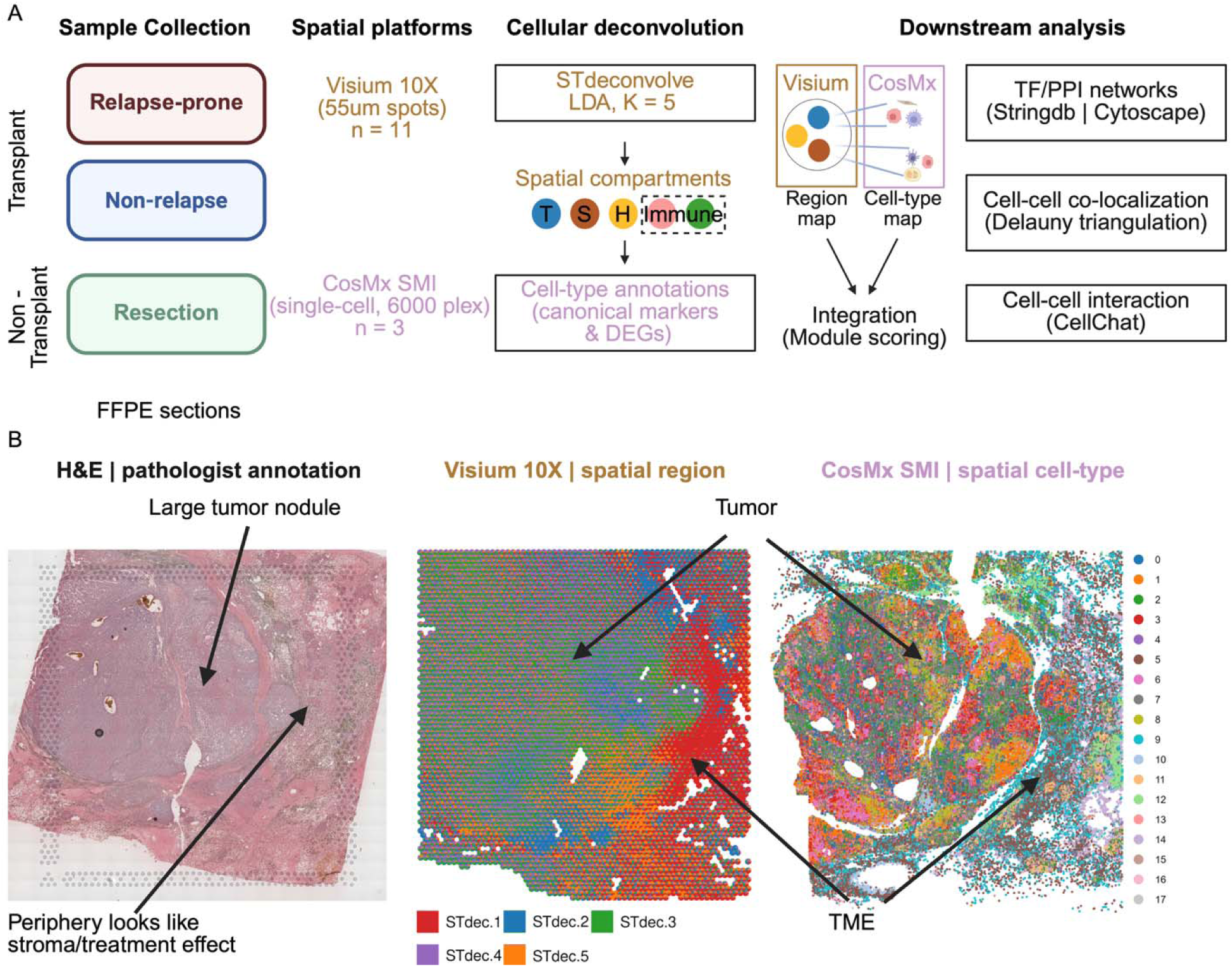
Integrative spatial transcriptomics workflow and cross-platform mapping of tumor architecture in relapse-prone unresectable hepatoblastoma. (A) Schematic overview of the study design. FFPE tumor sections from relapse-prone and non-relapsing unresectable HB, and HB cured with resection were profiled using Visium 10X (*n* = 11) and CosMx SMI (*n* = 3). Reference-free deconvolution of Visium data using STdeconvolve (LDA, *K* = 5) identified spatial compartments/regions, including tumor (T), stroma (S), hepatocyte-like (H), and immune-associated regions. Cell-type annotations were refined using canonical markers and differentially expressed genes (DEGs). Integration of Visium region-level and CosMx single-cell maps was performed via module scoring. Downstream analyses included cell-cell communication (CellChat), spatial co-localization (Delauny triangulation), and transcription factor/protein-protein interaction (TF/PPI) network analysis (Stringdb/Cytoscape). (B) Representative spatial correspondence across modalities. Left: H&E-stained section with pathologist annotation highlighting a large tumor nodule and peripheral stroma/treatment-associated regions. Middle: Visium-derived spatial compartments (STdec clusters) delineating tumor and surrounding microenvironment. Right: CosMx single-cell map showing high-resolution cell-type distribution, distinguishing tumor and TME compartments. Together, these data demonstrate concordance between histopathology, region-level transcriptomics, and single-cell spatial profiling.

Following quality control filtering, 32,960 features across 49,505 spatial spots from Visium and 6,175 features across 348,350 segmented cells from CosMx were retained for downstream analyses **(Fig. 2A-B)**. Each Visium spot (55 um diameter) captures transcripts from multiple cells (typically 5-50), necessitating deconvolution to infer cellular composition. To address this, we applied a reference-free deconvolution algorithm STdeconvolve and defined cell types (STdec) based on highly overdispersed genes (Methods; **Fig. 2C**). In parallel, unsupervised clustering identified transcriptional profiles independent of deconvolution **(Fig. 2D)**. Integration of these approaches with histological annotations enabled assignment of major spatial compartments, including tumor, stroma, hepatocyte, and immune-enriched regions (Methods; **Fig. 2E** and **Supplementary Fig. S1-2A** and **Table S4**). To link these region-level programs to single-cell resolution, we projected them onto CosMx data using module scoring, enabling resolution of the cellular state underlying each spatial compartment **(Supplementary Table S5)**. This integration establishes a multi-scale framework linking regional transcriptional programs with single-cell spatial organization.

**Fig. 2.**
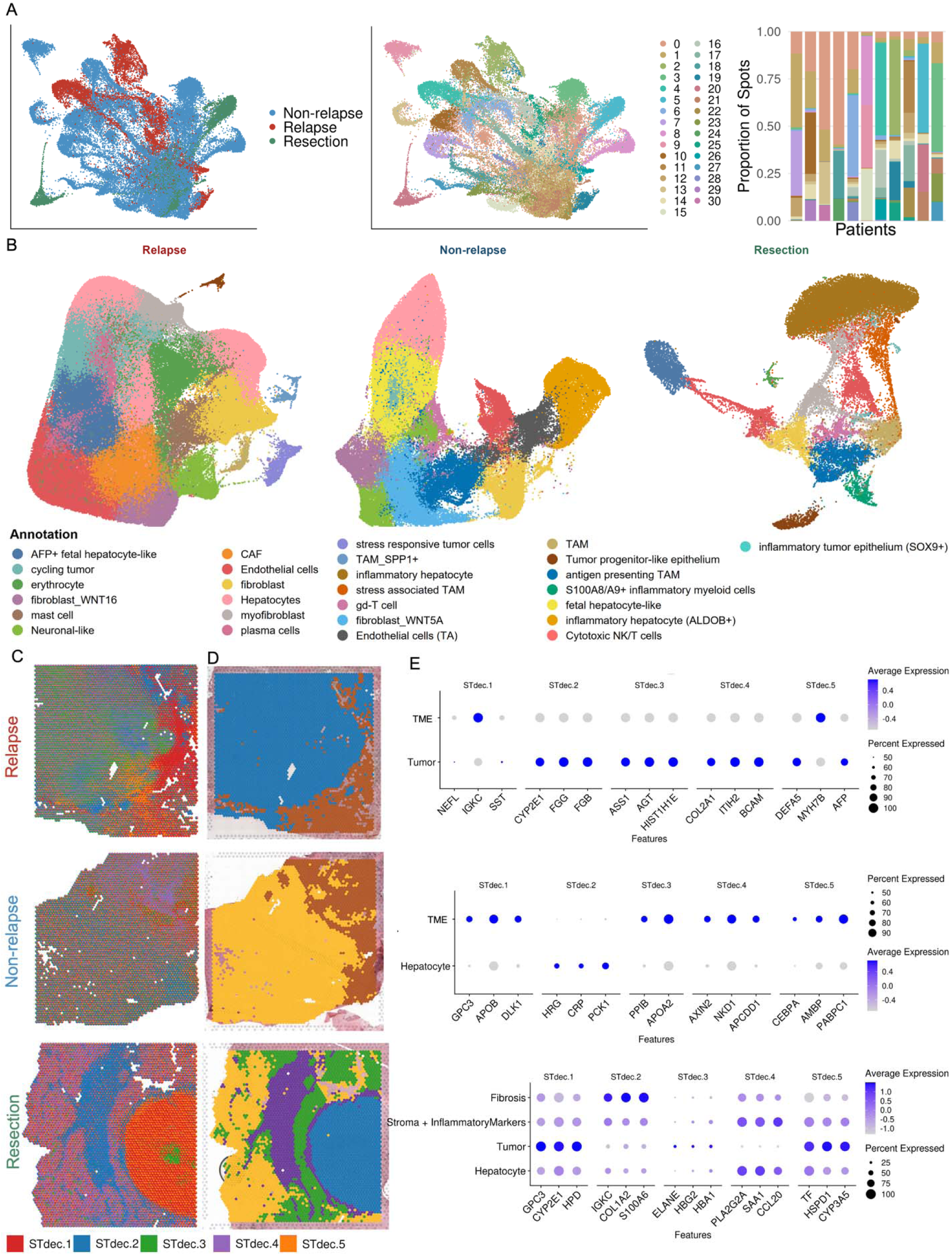
Spatially resolved tumor ecosystems across hepatoblastoma clinical groups. **(A)** Integrated low-dimensional embedding of Visium spots across all samples colored by clinical group (left; relapse, non-relapse, resection) and Seurat clusters (middle), revealing shared and distinct transcriptional programs. Right: stacked bar plot showing the proportion of Seurat clusters per patient, highlighting inter-patient heterogeneity and differential cluster enrichment across clinical groups. **(B)** High-resolution cell-state landscapes derived from CosMx SMI for representative samples from each clinical group (relapse, non-relapse, resection). Cells are annotated based on canonical markers and DEGs, capturing tumor, stromal, immune, hepatocyte-associated states. **(C)** Visium spatial maps of STdec topics (*K* = 5, STdec1 through STdec5) for representative sections from relapse, non-relapse, and resection samples, illustrating region-level compartmentalization of tumor and microenvironmental programs. **(D)** Corresponding region-level annotations derived from Seurat cluster profiling and marker-based interpretation, delineating tumor, stroma, hepatocyte-like, and inflammatory compartments across clinical groups. **(E)** Dot plots showing top 3 marker gene enrichment across STdeconvolve topics (*K* = 5) for each clinical group. Dot size represents the percentage of spots expressing each gene, and color indicates average expression. These profiles define the biological identity of spatial compartments, including tumor-associated programs, hepatocyte-like states, fibrosis/stromal signatures, and inflammatory or immune-associated features.

### Spatial architecture differs between relapse-prone, non-relapse, and resection hepatoblastoma samples

Relapse-prone tumors exhibit a TME-dominated and spatially coupled architecture. Mapping CosMx-derived cellular states to Visium-defined regions revealed marked differences in spatial organization across clinical groups **(Fig. 3A** and **Supplementary Fig. S2B-D)**. Importantly, this integration enabled resolution of cellular states not captured by STdec markers, refining the cellular composition of spatial regions. To derive mechanistic insights into the regulatory drivers of region-specific transcriptional programs, we integrated differential expression with protein-protein interaction (PPI) and TF-target network analyses.

**Fig. 3.**
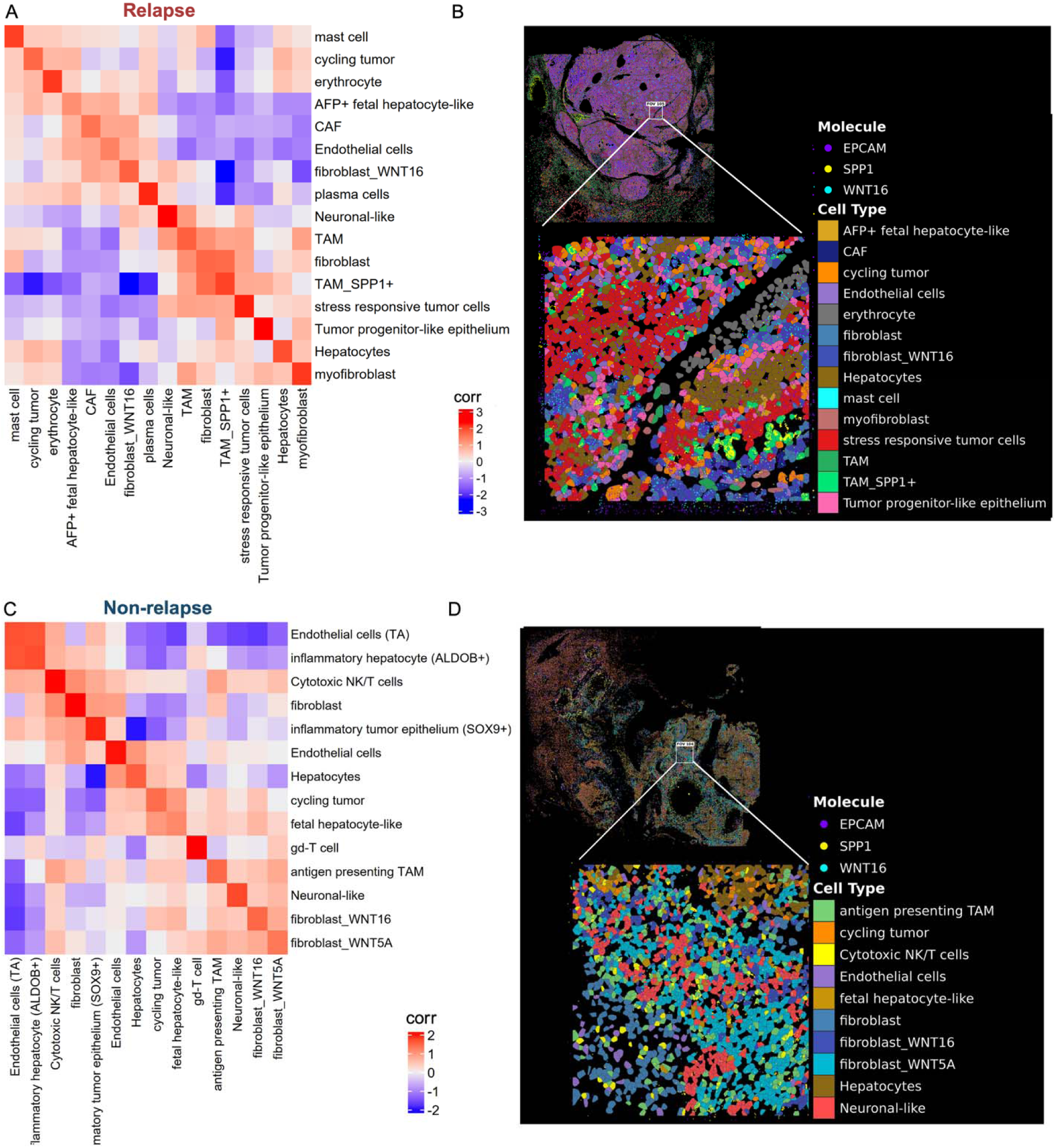
Spatial neighborhood enrichment analysis identifies cell-state co-localization patterns across hepatoblastoma clinical groups. (A, **C)** Heatmaps showing pairwise cell proximity enrichment scores computed from CosMx single-cell spatial data using Delaunay triangulation and permutation-based testing (*n* = 1000 simulations with FDR adjustment) for **(A)** relapse **(C)** non-relapse. Rows and columns represent annotated cell states. Positive values (red) denote cell-state pairs found in spatial proximity more often than expected by chance, whereas negative values (blue) denote spatial separation. **(B, D)** CosMx spatial maps of hepatoblastoma clinical groups. Top: whole-tissue overview of **(B)** relapse-prone, and **(D)** non-relapse, with EPCAM (violet), SPP1 (yellow), and WNT16 (cyan) transcripts overlaid on annotated cell types; white box marks the zoomed field of view (FOV) (Bottom). **(B)** In relapse, the zoomed FOV shows tumor cells intermixed with TME, with a discrete TAM_SPP1 –fibroblast niche, consistent with a macrophage-fibroblast regulatory circuit driving tumor plasticity. **(D)** In non-relapse, the zoomed FOV reveals fibroblast_WNT5A /WNT16 populations intermixed with antigen-presenting TAMs, with a discrete cytotoxic NK/T cell–fibroblast niche, consistent with a compartmentalized tumor-immune neighborhood.

Within post-chemotherapy unresectable samples, relapse-prone tumors exhibited high tumor cell-type and-state heterogeneity, with tumor regions composed of progenitor-like, stress-responsive, *AFP* fetal hepatocyte-like, and cycling tumor cells embedded within a microenvironment enriched for *SPP1* tumor-associated macrophages (TAMs), fibroblasts, hepatocytes, plasma cells, and myofibroblasts. Furthermore, TME regions were enriched for plasma cells, myofibroblasts, mast cells, erythrocytes, and hepatocyte-like populations, with fewer tumor cell states **(Supplementary Fig. S2B)**. CosMx further resolved additional populations including neuronal-like cells (TME regions), endothelial cells (tumor and TME regions), fibroblast_*WNT16* populations, and CAFs within tumor regions that were not identified by STdec markers, indicating unresolved layers of spatial heterogeneity. Region-specific differential expression analysis from Visium data showed that tumor regions were enriched for proliferative, MAPK, and Wnt signaling programs, whereas TME regions were enriched for extracellular matrix remodeling, interferon response, and immune regulatory pathways **(Supplementary Fig. S3A)**. Collectively, these features are consistent with aggressive tumor behavior where a TME-dominated regulatory environment supports tumor plasticity and heterogeneity.

In contrast, non-relapse tumors display a more distributed and immune-engaged spatial organization. These tumors displayed a more integrated and less tumor-dominated architecture, characterized by prominent hepatocyte-associated programs and broader immune engagement. Tumor regions contained cycling tumor cells, hepatocyte-like states, fetal hepatocyte-like populations, neuronal-like cells, and fibroblast populations including *WNT16* and *WNT5A* fibroblasts. While hepatocyte regions were enriched for canonical hepatocytes, inflammatory hepatocytes (*ALDOB*), cycling tumor cells, and fetal hepatocyte-like populations. Several immune and stromal populations not captured by STdec markers including antigen-presenting TAMs, fibroblasts, endothelial cells, and cytotoxic NK/T cells were distributed across regions rather than confined to discrete regions. Additional cell-types and-states, including inflammatory tumor cells (*SOX9*) and γδ T cells, were specifically associated with hepatocyte regions and were likewise not identified by STdec markers **(Supplementary Fig. S2C)**. The coexistence of tumor and hepatocyte transcriptional programs within these regions suggests active tumor remodeling toward hepatocyte-like states. Region-specific differential expression analysis from Visium data showed that tumor regions were enriched for vascular-ECM signaling, metabolic and stress-response pathways, and signal transduction programs, whereas TME regions exhibited enrichment of fibroblast/myeloid signaling, immune regulation, and interferon response **(Supplementary Fig. S3B)**. Together, these findings support a model of distributed tumor-immune crosstalk and hepatocyte-associated tumor remodeling, consistent with a more immune-engaged and constrained tumor ecosystem.

Resectable tumors exhibit a more structured and stabilized tumor microenvironment. In contrast to unresectable samples, these tumors represent a spatially organized and constrained architecture, although CosMx profiling depth was limited by technical factors. Resectable samples showed reduced tumor cell-type and state heterogeneity relative to relapse-prone tumors and increased immune diversity compared to non-relapse tumors. Tumor regions contained *AFP* fetal hepatocyte-like tumor cells, while fibrotic regions were enriched for fibroblasts and plasma cells. Hepatocyte and TME regions contained inflammatory hepatocytes, myofibroblasts, stress-associated TAMs, S100A8/A9 inflammatory myeloid cells, and erythrocytes. Furthermore, antigen-presenting TAMs and endothelial cells were distributed across multiple regions, suggesting active immune response. Additional cell populations including TAMs, tumor progenitor-like epithelial cells, γδ T cells, and cycling tumor cells were primarily associated with hepatocyte and stroma regions but were not captured by STdec markers **(Supplementary Fig. S2D)**. Region-specific differential expression analysis from Visium data showed that tumor regions were enriched for epithelial proliferation, lipid and cholesterol biosynthesis, and Wnt/β-catenin signaling, alongside regulators of epithelial identity and transcriptional plasticity, consistent with a metabolically active, Wnt-driven epithelial state with transcriptional reprogramming and adaptive plasticity. In contrast, TME regions were enriched for macrophage differentiation, interferon response, and tissue remodeling programs **(Supplementary Fig. S3C)**. Together, these findings suggest a tumor ecosystem in which cellular adaptation is driven primarily by transcriptional and microenvironmental programs rather than genomic instability. Collectively, these findings define three distinct spatial organizational states: a TME-dominated, centrally regulated architecture in relapse-prone tumors, a distributed and immune-engaged architecture in non-relapse tumors, and a stabilized, constrained architecture in resection samples. A summary of the major spatial, cellular, and signaling features associated with each clinical group is provided in **Supplementary Table S6**.

### Spatial proximity analysis defines functionally distinct tumor neighborhoods across hepatoblastoma clinical groups

To define higher-order spatial organization beyond cell-type composition, we analyzed cell-cell co-localization using Delaunay triangulation-based networks. This approach identified recurrent cellular neighborhoods across clinical groups, defined by spatial proximity of specific cell types. To characterize the functional programs associated with these neighborhoods, we performed pathway enrichment analysis on differentially expressed genes (FDR ≤ 0.01, log2FC ≥ 0.05) of co-localized cell types.

Relapse-prone tumors are organized around tightly coupled tumor-TME-immune architectures. In these tumor regions, spatial co-localization analysis revealed a dominant cellular neighborhood in which tumor cells (progenitor-like and stress-responsive cell states), fibroblasts, myofibroblasts, and macrophages (*SPP1* TAMs and TAMs) co-localized, forming a TME-dominated regulatory environment (**Fig. 3A).** This neighborhood was spatially validated by the representative in situ CosMx FOV, which showed a discrete TAM_SPP1 –fibroblast niche embedded within a region where tumor cells intermixed with TME **(Fig. 3B).** Within this neighborhood, tumor cells were enriched for hypoxia-response and metabolic reprograming, consistent with adaptive stress states, while TME populations contributed to extracellular matrix remodeling, migration, and Wnt/MAPK activity. Macrophages exhibited inflammatory and antigen presentation signatures, and co-localized broadly with multiple cell-types and states, consistent with a hub-like regulatory role in coordinating TME-tumor interactions. In TME regions, co-localization of myofibroblasts, neuronal-like, and macrophages defined a secondary neighborhood enriched for extracellular matrix organization and inflammatory signaling, indicating compartmentalized TME remodeling supporting tumor progression. These spatially defined neighborhoods further resolve the cellular basis of the TME-dominated regulatory environment. Collectively, these findings support a model in which relapse-prone tumors are organized around a macrophage-fibroblast regulatory circuit that promotes hypoxia-adaptive tumor plasticity and therapy-resistant states **(Fig. 3A)**.

Non-relapse tumors exhibit compartmentalized and functionally distinct cellular neighborhoods. These tumors revealed multiple co-existing neighborhoods, reflecting localized tumor-immune crosstalk and compartmentalized microenvironmental regulation. In tumor regions, two functionally distinct neighborhoods were observed: (i) cycling tumor cell states co-localized with fetal hepatocyte-like states, forming a metabolically regulated cellular neighborhood, and (ii) cytotoxic NK/T cells co-localized with fibroblast, defining a pro-inflammatory, ECM remodeling neighborhood consistent with immune-engagement. Similarly, TME regions were organized into distinct cellular neighborhoods. Endothelial cells co-localized with inflammatory hepatocyte, forming an angiogenic and metabolically active neighborhood, while fibroblast co-localized with inflammatory epithelium and cytotoxic NK/T cells, exhibiting stress-response and inflammatory signaling **(Fig. 3C)**. Consistent with this, the representative in situ CosMx FOV showed fibroblast_WNT5A and fibroblast_WNT16 populations intermixed with antigen-presenting TAMs, alongside a discrete cytotoxic NK/T cell–fibroblast niche within the compartmentalized tumor-immune neighborhood **(Fig. 3D)**. This spatial organization parallels the underlying transcriptional programs and highlights localized tumor-immune crosstalk and compartmentalized microenvironmental regulation.

Resection tumors displayed more uniform and recurrent spatial interactions. In contrast to unresectable tumors, resection tumors represented a distinct spatial organization characterized by reduced tumor heterogeneity and increased uniformity of cellular interactions. Resection samples displayed homogeneous and recurrent cellular neighborhoods, characterized by consistent co-localization of TME and immune populations across tissue regions. Dominant cellular neighborhoods involved fibroblasts, antigen-presenting macrophages, and tumor progenitor-like cells, alongside adaptive immune populations, indicating a structured and stabilized tumor-microenvironment architecture **(Supplementary Fig. S4A)**. This organization was corroborated by the representative in situ CosMx FOV, which showed a discrete SPP1 tumor progenitor plasma cell niche, consistent with the fibroblast-antigen-presenting TAM-tumor progenitor cluster identified in **Supplementary Fig. S4A**, embedded within a separately admixed inflammatory hepatocyte TAM myofibroblast background (**Supplementary Fig. S4B)**. This recurrent pattern reflects the coordinated transcriptional and microenvironmental programs observed at the regional level, indicating a spatially structured and functionally coordinated TME-immune network with reduced heterogeneity of tumor-associated cell states.

Distinct cell-cell signaling programs underlie these spatially organized cellular neighborhoods. To define the functional communication underlying these cellular neighborhoods, we performed ligand-receptor interaction analysis. Ligand-receptor analysis revealed distinct dominant signaling programs across clinical groups, with Midkine signaling enriched in relapse-prone tumors, primarily involving interactions between endothelial cells, CAFs, and *AFP* fetal hepatocyte-like tumor cell states, whereas collagen-mediated signaling predominated in non-relapse and resection samples, involving fibroblasts, myofibroblasts, and immune populations **(Fig. 4A)**. Collectively, these findings demonstrate that spatial proximity and functional communication represent complementary layers of tumor organization, together defining the regulatory architecture of hepatoblastoma across distinct clinical contexts.

**Fig. 4.**
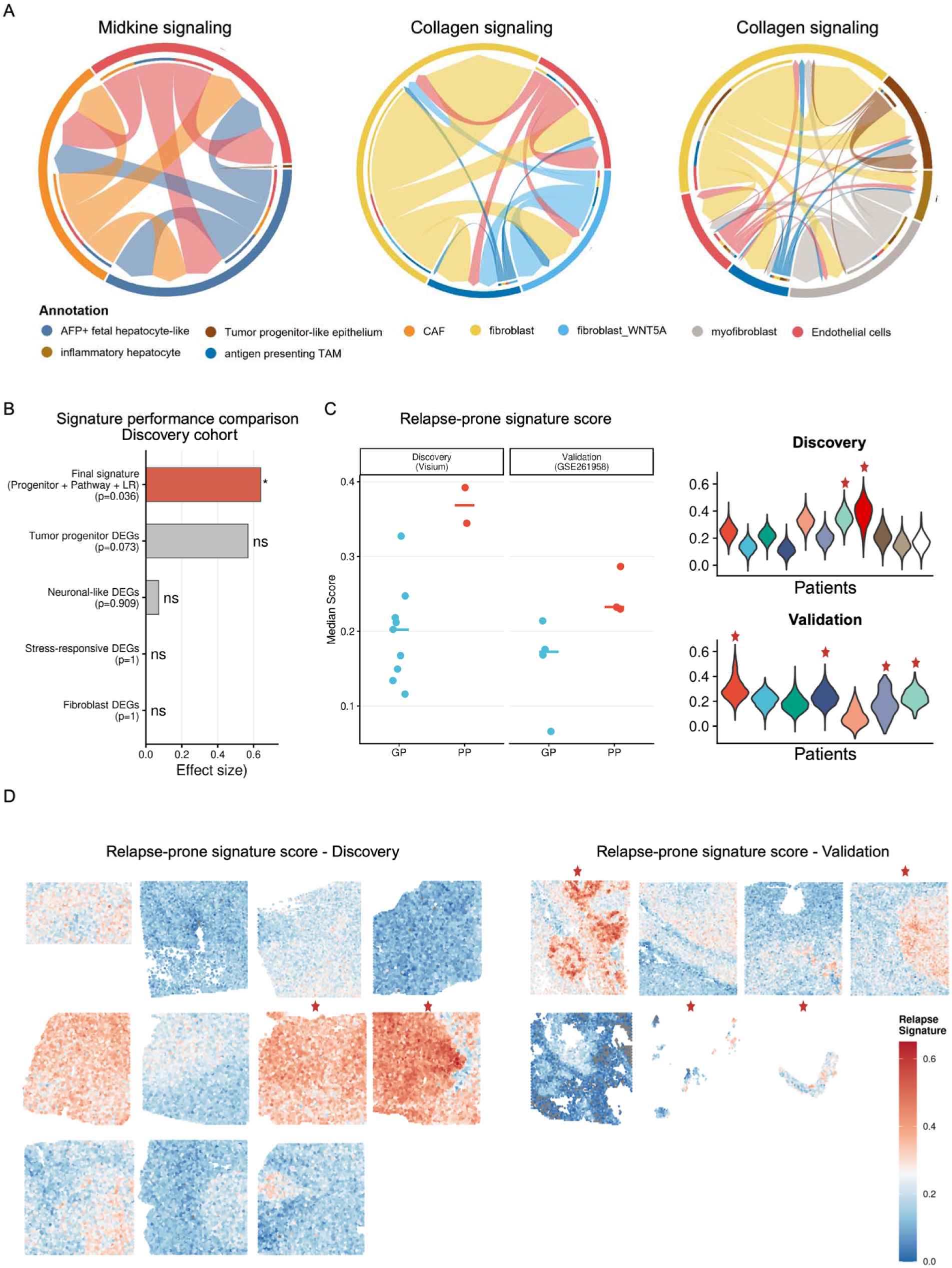
Spatially resolved signaling programs define relapse-associated tumor architecture and prognostic signatures. **(A)** Ligand-receptor interaction analysis of spatially defined cellular neighborhoods reveals distinct dominant signaling programs across clinical groups. Relapse-prone tumors are characterized by enriched Midkine signaling, primarily involving interactions between endothelial cells, cancer-associated fibroblasts (CAFs), and *AFP* fetal hepatocyte-like and tumor progenitor-like epithelial states. In contrast, non-relapse and resection samples exhibit dominant collagen-mediated signaling, driven by fibroblast, myofibroblast, and immune-associated interactions. Networks represent aggregated interaction strengths inferred using CellChat. **(B)** Comparative performance of candidate gene signature for relapse prognostic in the discovery cohort. The final composite signature (tumor progenitor + pathway + LR features) demonstrates improved effect size relative to individual gene sets, indicating that integration of spatial and signaling features enhances prognostic power. **(C)** Relapse-prone signature scores across discovery (Visium) and validation cohorts. Relapse-associated samples exhibited significantly higher signature scores compared to non-relapse samples, demonstrating robustness across independent datasets. Violin plots show per-patient distribution, red asterisks indicate relapse samples. **(D)** Spatial distribution of relapse-prone signature scores tissue sections in discovery and validation cohorts. High-scoring regions localize to distinct spatial niches, red asterisks indicate relapse samples.

**Fig. 5.**
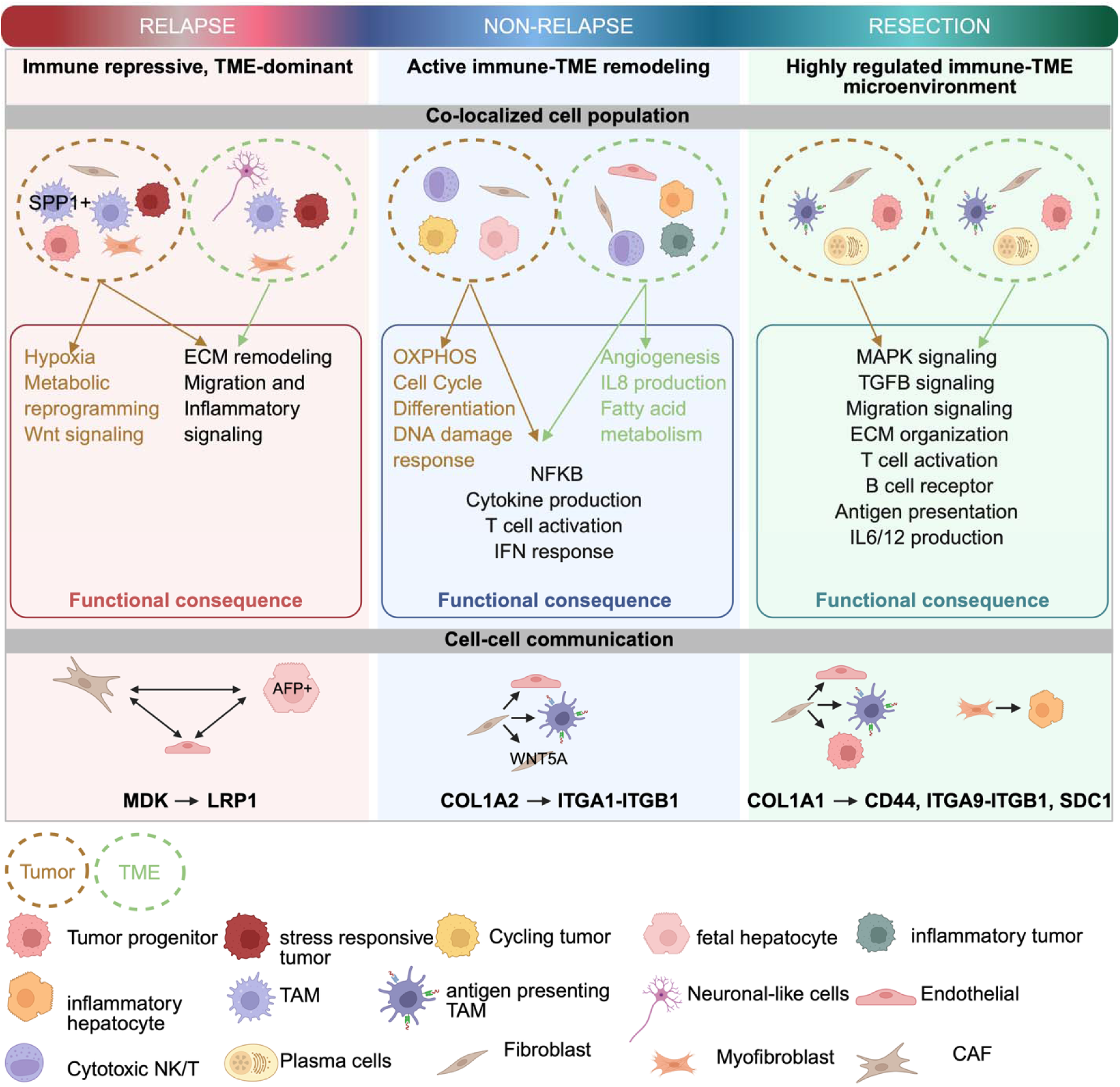
Distinct spatial architectures and signaling programs define relapse-associated and non-relapse hepatoblastoma microenvironments. Schematic representation of spatially resolved tumor ecosystems across clinical groups, integrating cell composition, functional programs, and dominant cell-cell communication pathways. Relapse-prone tumors exhibit an immune-repressive, TME-dominant microenvironment characterized by co-localization of tumor progenitor-like cells with TME populations, including CAFs and TAMs. These architectures are associated with hypoxia, metabolic reprogramming and WNT signaling, and driven by Midkine mediated interactions, suggesting a pro-tumorigenic and therapy-resistant state. Non-relapse tumors display active immune-TME remodeling, with spatially coordinated interactions between tumor, immune, and stromal populations. These regions are enriched for oxidative phosphorylation, cell cycle activity, and inflammatory signaling, including NF-kB activation and cytokine production. Collagen-mediated signaling contributes to this condition. Resection samples exhibit a highly regulated immune-TME microenvironment characterized by organized interactions between immune and TME compartments. These regions are enriched for MAPK and TGFb signaling, ECM organization, and adaptive immune responses, including T-and B-cell activation and antigen presentation. Collagen-driven signaling underlies these changes.

### Validation of relapse-associated tumor progenitor signature across independent cohort

We next assessed whether relapse-associated spatial niches translate into a clinically relevant gene signature. To validate this, we derived a gene signature capturing the tumor-intrinsic transcriptional state of the relapse-associated microenvironment. To identify the optimal cellular source, we first compared the discriminative performance of DEG sets from each tumor niche relapse-associated cell types in the discovery cohort. Tumor progenitor-like DEGs showed the strongest discrimination (effect size = 0.569, reflecting magnitude of separation between groups, *p* = 0.073), while neuronal-like, fibroblast, and stress-responsive DEG sets showed no signal (effect size ≤ 0.071, *p* ≥ 0.909; **Fig. 4B**), indicating tumor progenitor-like cells as the signature source. Consistent with this, liver progenitor populations have been reported to expand following neoadjuvant chemotherapy and are enriched in relapse and metastatic settings (30).

The final signature was constructed by integrating tumor progenitor DEGs, pathway-enriched genes from EnrichR analysis, and ligand-receptor interaction genes from high-confidence cell-cell communication analysis (*p* < 0.01, top 90th percentile), particularly among CAFs and AFP fetal hepatocyte-like cell states. This integration improved discrimination beyond DEGs alone, yielding a 68-gene signature with effect size 0.640 (*p* = 0.036; **Fig. 4A**). Six genes including *FGFR*, *FGFR4*, *CXCL12*, *AGTR1*, *LRP1*, and *LAMC1* were supported by both differential expression and active intercellular signaling, representing a high-confidence core. In the discovery cohort (*n* = 11, Visium), the signature was significantly enriched in relapse versus non-relapse patients (*p* = 0.036, median effect = 0.163), with both relapse patients scoring above the non-relapse median **(Fig. 4C)**. Spatial visualization revealed focal enrichment at tumor-TME boundaries **(Fig. 4D)**. Applied to an independent external cohort (31) (GSE261958, CytAssist FFPE, *n* = 7), patients with poor outcome scored consistently higher than survivors (*p* = 0.057, median effect = 0.063), supporting directional consistency across independent platforms and cohorts **(Fig. 4C)**.

## Discussion

Therapeutic response in hepatoblastoma is reflected in the spatial architecture of the tumor ecosystem rather than cellular composition alone. Using integrated regional and single-cell spatial transcriptomics, we demonstrate that post-chemotherapy hepatoblastoma is reorganized into distinct regulatory architectures that differ systematically across clinical outcomes. These architectures represent therapy-remodeled tumor ecosystems in which tumor plasticity, TME organization, and immune engagement are coordinately restructured. These patterns can be broadly categorized into constrained (resection), distributed (non-relapse), and centralized (relapse) architectures **Supplementary Table S6**.

Myeloid cells are known to play a central role in hepatoblastoma and other solid tumors, particularly in shaping tumor-stroma interactions and therapeutic response (13,30,32). A defining feature of the relapse-prone microenvironment in our study is the spatial enrichment of *SPP1* TAMs within a TME-dominated neighborhood co-localizing with CAFs and *AFP* fetal hepatocyte-like tumor cells. *SPP1* TAMs have been broadly associated with poor prognosis and linked to hypoxia, TGFb signaling, and immune exclusion, with CXCL9:SPP1 macrophage polarization emerging as a key determinant of outcome across cancers (33–35). However, prior studies have primarily focused on macrophage abundance and transcriptional state. Here, we show that *SPP1* TAMs are centrally positioned within TME, where they co-localize with fibroblasts and tumor cells and coordinate their interactions following chemotherapy. Within this architecture, TME populations, including stromal and endothelial populations show strong interactions with *AFP+* tumor cell states, driven in part by Midkine signaling via MDK-LRP1 ligand-receptor interaction, consistent with coordinated regulation of tumor plasticity and immune evasion (31). Beyond Midkine signaling, the relapse-associated transcriptional program is enriched for pathways linked to chemokine signaling, extracellular matrix remodeling, angiogenesis, and receptor tyrosine kinase activation, as reflected by genes such as *CXCL12*, *LAMC1*, *AGTR1*, *LRP1*, and *FGFR4*. Notably, *FGFR4* has been associated with proliferative, chemotherapy-resistant hepatoblastoma tumor cell states and is enhanced through fibroblast-tumor signaling interactions, supporting a role for FGF-FGFR signaling as a stromal-reinforced growth axis in this spatial neighborhood (15,36–39). These findings align with prior work demonstrating roles for *SPP1* TAMs in promoting tumor stemness, metabolic dysregulation, and immune suppression (40), as well as TAM-CAF interactions in desmoplastic remodeling (41). Importantly, our data extend these observations to hepatoblastoma and demonstrate that these processes are spatially organized, highlighting a coordinated stromal-immune regulatory architecture associated with therapy-resistant tumor states.

Hepatoblastoma prognosis is closely linked to its inherent histological heterogeneity, with specific epithelial cell states—including high-risk small cell undifferentiated populations— defining clinical behavior (42,43). Correspondingly, bulk transcriptomic analysis reveals that epithelial gene expression programs drive tumor risk stratification (9). Our study defines a distinct, context-dependent landscape of hepatoblastoma cell states: relapse-prone tumors are enriched for progenitor-like, cycling, and stress-responsive cells, while non-relapse cases exhibit more differentiated, inflammatory epithelial states. Furthermore, resection samples show reduced heterogeneity with a predominance of fetal hepatocyte-like and progenitor-like states. Spatial analysis reveals that the tumor microenvironment (TME), specifically immune and stromal components, actively shapes these tumor cell compositions. Non-relapse tumors maintain a distinct differentiation hierarchy, comprising inflammatory hepatocyte-like cells and inflammatory tumor progenitors. While previous studies show that treatment shifts hepatoblastoma toward hepatocytic differentiation, specifically enriching HNF1A/HNF4A programs (30), our data indicates that these inflammatory hepatocyte-like states also harbor mature signatures alongside acute-phase, stress-adaptive responses. These states localize to regions with treatment-associated necrosis and fibrosis **(Supplementary Table S2)**. Conversely, inflammatory tumor progenitor-like populations retain developmental signaling programs, such as Wnt and Notch pathway activity, highlighting the persistence of regulated tumor plasticity. Spatially, these epithelial states exist within microenvironments enriched for antigen-presenting macrophages, cytotoxic lymphocytes, and stromal populations.

Limitations of this study include a small, predominantly post-treatment sample size, which may bias results toward chemo-resistant states and underrepresent pre-treatment heterogeneity. However, the study focuses on the highest-risk tumors because most patients underwent liver transplantation for unresectable disease and were maintained on post-transplant immunosuppression at the time of relapse. Immunosuppression inhibits tumor immune surveillance, creating conditions uniquely permissive to residual disease proliferation. Post-transplant HB relapses are known to be rapid and fatal within a year or two. No prior spatial transcriptomic study of hepatoblastoma has been conducted in this clinical context, and the fact that our spatial signatures distinguished relapsing from non-relapsing patients within this uniformly immunosuppressed cohort strengthens the argument that tumor-TME architecture carries prognostic information independent of immune status. While our spatial transcriptomics approach offers multi-scale insights, further studies with large cohorts and functional validation will be necessary to confirm the causal role of these architectures in driving therapeutic resistance. Looking ahead, advanced single-nucleus profiling will allow for broader analysis. In conclusion, we show that post-chemotherapy hepatoblastoma is organized into distinct spatial regulatory states link to clinical outcome. These findings position spatially organized tumor-microenvironment interactions as central determinants of therapeutic response and highlight microenvironmental signaling networks as targets of overcoming treatment resistance.

## Acknowledgments

The clinical work was supported by Raizman-Haney Endowment–RS; Hillman Foundation of Pittsburgh; Szalay Family Foundation; Janet Family; Pediatric Surgery Scientific Program to RS; the systems biology work was supported by NIH grants, OT2-OD030544 OT2-OD03643, and a Wellcome Leap grant to SS.

## Data and Code Availability Statement

Visium data are available from NCBI GEO (GSEXXXX), and CosMx SMI data are provided at [GSEXXXX]. Analysis code is available at https://github.com/zahramesri/Hepatoblastoma_CosMx-Visium.

